# Dynein dysfunction prevents maintenance of high concentrations of slow axonal transport cargos at the axon terminal: a computational study

**DOI:** 10.1101/2022.06.19.496644

**Authors:** Ivan A. Kuznetsov, Andrey V. Kuznetsov

## Abstract

Here we report computational studies of bidirectional transport in an axon, specifically focusing on predictions when the retrograde motor becomes dysfunctional. We are motivated by reports that mutations in dynein-encoding genes can cause diseases associated with peripheral motor and sensory neurons, such as type 2O Charcot-Marie-Tooth disease. We use two different models to simulate bidirectional transport in an axon: an anterograde-retrograde model, which neglects passive transport by diffusion in the cytosol, and a full slow transport model, which includes passive transport by diffusion in the cytosol. As dynein is a retrograde motor, dysfunction should not directly influence anterograde transport. However, our modeling results unexpectedly predict that slow axonal transport fails to transport cargos against their concentration gradient without dynein. The reason is the lack of a physical mechanism for the reverse information flow from the axon terminal, which is required so that the cargo concentration at the terminal could influence the cargo concentration distribution in the axon. Mathematically speaking, to achieve a prescribed concentration at the terminal, equations governing cargo transport must allow for the imposition of a boundary condition postulating the cargo concentration at the terminal. Perturbation analysis for the case when the retrograde motor velocity becomes close to zero predicts uniform cargo distributions along the axon. The obtained results explain why slow axonal transport must be bidirectional to allow for the maintenance of concentration gradients along the axon length. Our result is limited to small cargo diffusivity, which is a reasonable assumption for many slow axonal transport cargos (such as cytosolic and cytoskeletal proteins, neurofilaments, actin, and microtubules) which are transported as large multiprotein complexes or polymers.

## 1. Introduction

Neurons utilize a complicated “railway” system, composed of microtubule (MT) tracks and molecular motors that move on these tracks, to transport large proteins and vesicles across large distances. The motors themselves belong to the kinesin family, which perform anterograde transport, or the dynein family, which perform retrograde transport [1].

Motor protein dysfunction plays a critical role in many neurodegenerative diseases [2–4]. Dynein mutations have been implicated in many cases of malformations of cortical development, spinal muscular atrophy with lower extremity dominance, and congenital muscular dystrophy. Another example is Charcot-Marie-Tooth (CMT) disease, a disorder of peripheral motor and sensory neurons [5]. CMT type 2O disease is caused by a mutation in gene DYNC1H1 that encodes the dynein heavy chain [5–8]. Although the involvement of molecular motors in CMT pathogenesis is not surprising – long sensory and motor neurons require a large anterograde flux of various cargos from the soma to support their remote terminals [9] – an intriguing question is why degeneration of these neurons is associated with dynein dysfunction. Previous hypotheses as to why neurons can be negatively affected by dynein dysfunction include an inability to deliver dysfunctional organelles/proteins to the soma for degradation or a failure of retrograde signaling [10]. Bidirectional transport may be also needed to correct errors if cargo is delivered to a wrong location [11].

Here, we consider another possibility that is associated with our hypothesis that dynein-driven transport is necessary for cargo transport against its concentration gradient. Consider a mathematical argument: when cargo diffusivity is small, a continuum model of anterograde transport is described by a first-order differential equation, which allows for the imposition of a boundary condition only at the axon hillock. The equation does not allow for the imposition of the second boundary condition (prescribing a higher cargo concentration) at the axon tip. To simulate an increased cargo concentration at the axon tip, both anterograde and retrograde components are needed [12]. Ref. [12] used this argument to explain the presence of a retrograde component in slow axonal transport. Note that according to our hypothesis dynein-driven transport does not only serve for signaling to the soma to correctly modulate protein synthesis and to regulate forward kinesin transport. Rather, anterograde and retrograde transport are coupled at any location along the axon to form a physical system that allows specifying cargo concentrations at both ends of the axon, at the hillock and at the terminal.

In this paper, we attempt to answer the question of how dynein-driven transport can affect the transport of cargos toward the axon terminal. We will focus on slow axonal transport-b (SCb), which is used to transport ~200 proteins from the soma to the presynaptic terminal [13]. SCb transports cargo at an average velocity of 2-8 mm/day (0.023-0.093 μm/s), much less than the 1 μm/s observed for kinesin and dynein motors. This slower SCb transport velocity is explained by significant time cargos spend in the pausing state. Another interesting feature of SCb is that during the rapid phase of motion the cargos move bidirectionally with a bias toward anterograde motion [14,15]. An intriguing question raised in ref. [10] and by many other researchers is then: why is slow axonal transport bidirectional if cargoes just need to be transported to the axonal tip? Since molecular motors use ATP, bidirectional transport is much more energetically expensive than unidirectional transport. One possible explanation is that bidirectional cargo movements are caused by opposite polarity motors that are attached to the same cargo and that are in tug-of-war [16,17]. Here, we suggest that bidirectional transport is required to maintain an increasing cargo concentration along the length of the axon, in the situation when the cargo concentration is small at the soma and high at the presynaptic terminal. This explanation first reported in ref. [12] holds for the case when cargos are transported as polymers or large multiprotein complexes that have small diffusivity, which is the case for many SCa and SCb proteins.

We focus on one particular slow axonal transport cargo, α-synuclein (α-syn), mostly known for its involvement in Parkinson’s disease [18–23]. Since in a healthy neuron α-syn predominantly exists in the monomeric form [24], hereafter we denote α-syn monomer as α-syn.

Our model is a cargo-level model that simulates the behavior of cargos rather than the behavior of motors, as a specific cargo can be driven by several motors. We simulate dynein dysfunction as a decrease of dynein velocity. This is equivalent to specifying the velocity of retrograde cargos during the fast phase of their movement on MTs. Note that cargos can also pause when the motors that drive them temporarily disengage from MTs.

Our goal is to simulate how dynein dysfunction affects slow axonal transport of α-syn. Although we perform computations using α-syn parameters, we expect our results (the need for the retrograde component) to be generalizable to other slow axonal transport cargos.

## 2. Methods and models

### 2.1. A simplified case: axonal transport model that includes anterograde and retrograde motor-driven transport without diffusion and pausing

#### 2.1.1. General formulation of the anterograde-retrograde cargo transport model

A schematic representation of an axon is shown in Fig. 1. We start with the simplified model of bidirectional transport suggested in ref. [12]. Since long-distance transport of α-syn is propelled by kinesin and dynein motors [25], the simplified model includes only two kinetic states, which simulate anterograde (driven by kinesin) and retrograde (driven by dynein) cargos (Fig. 2a).

**Fig. 1.**
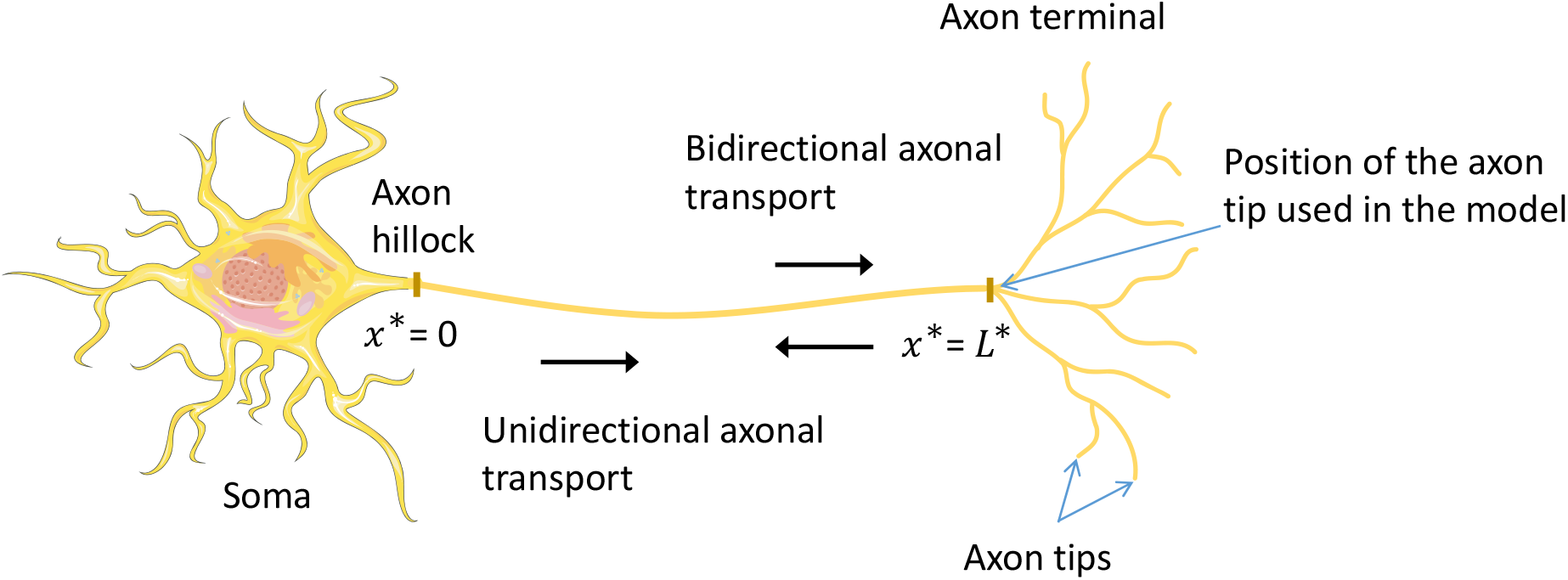
A diagram that defines the coordinate system adopted in the models. The *x*\*-coordinate defines distance from the axon hillock (*x*\* = 0) to the axon tip (*x*\* = *L*\*). Figure is generated with the aid of Servier Medical Art, licensed under a creative commons attribution 3.0 generic license, http://Smart.servier.com.

**Fig. 2.**
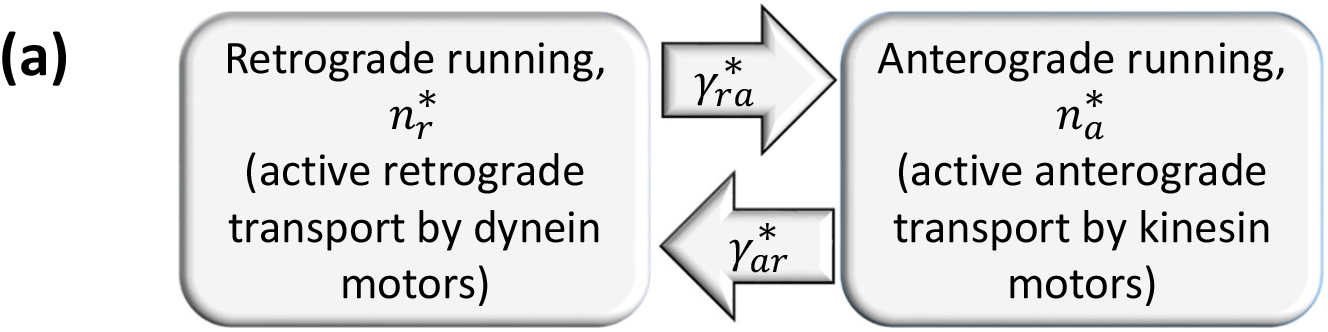
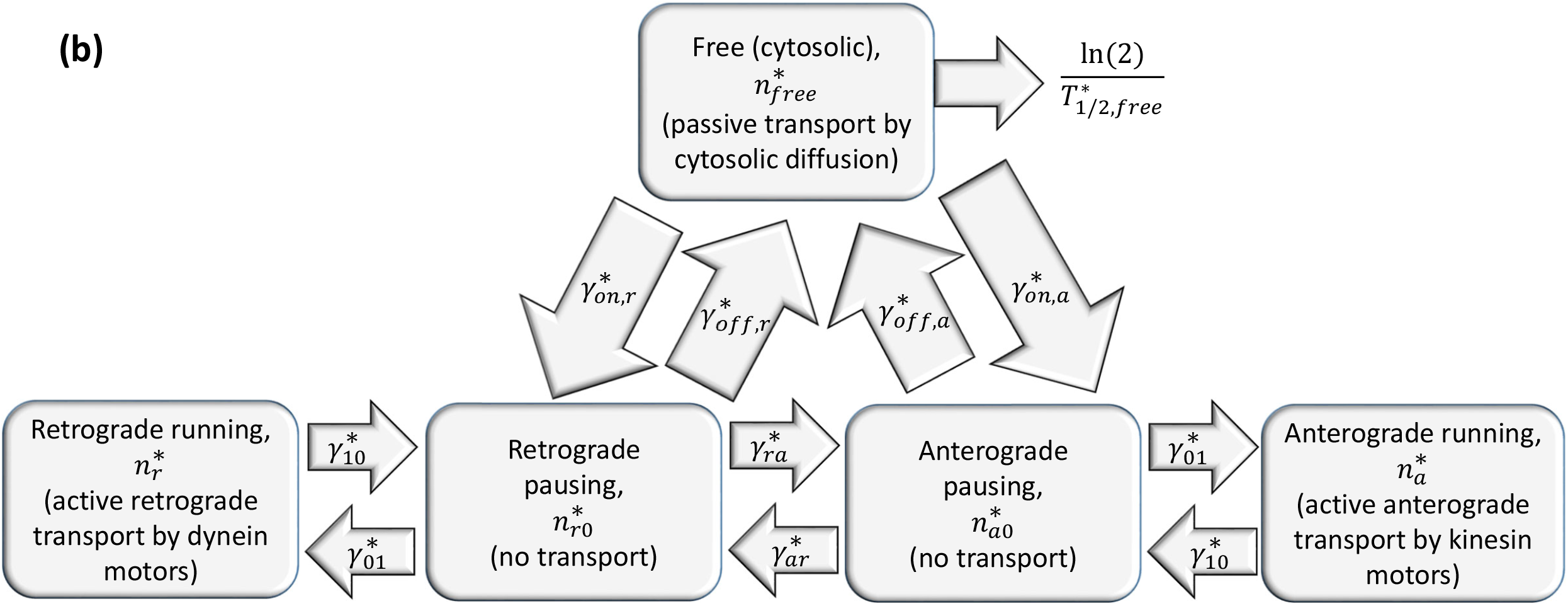
Kinetic diagrams that depict various kinetic states in two models of α-syn transport in the axon and transitions between these kinetic states. (a) A diagram for the model with anterograde and retrograde motor-driven kinetic states (without cargo diffusion and pausing states). (b) A diagram for the full slow axonal transport model. The diagram is based on the model of SCa transport of neurofilaments developed in ref. [26] with modifications to this model suggested in refs. [27,28] to extend this model to cytosolic proteins transported in SCb. The full slow axonal transport model also includes degradation of free α-syn due to its destruction in proteasomes.

For simulating long-range cargo transport in an axon, we used a quasi-steady-state approximation and neglected the time derivatives in the model equations. We also simulated axonal transport as one-dimensional and characterized cargo concentrations in all kinetic states by their linear number densities. We defined α-syn concentrations in various kinetic states in Table S1. Model parameters are defined in Tables S2 and S3. Table S3 defines kinetic constants, γ*s, used in the model to characterize α-syn transitions between different kinetic states, see Fig. 2a. Stating the conservation of α-syn in the anterograde state, characterized by concentration 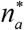 (Fig. 2a), we obtained the following equation:

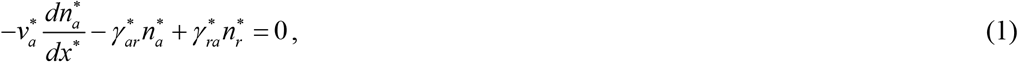

where asterisks denote dimensional quantities, *x*\* is the Cartesian coordinate that protrudes from the axon hillock (*x*\* = 0) to the axon tip (*x*\* = *L*\*). Sticky speaking, the positions of axon tips are shown in Fig. 1; however, we do not resolve complicated processes occurring in the terminal, our model is only valid between 0 ≤ *x*\* ≤ *L*\*. 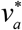 is the velocity of rapid motions of α-syn on MTs propelled by kinesin motors in slow axonal transport, and 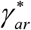 and 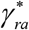 are kinetic constants defined in Fig. 2a. The first term on the left-hand side of Eq. (1) characterizes a change of the number of α-syn molecules in the control volume (CV) due to anterograde transport of α-syn, the second term characterizes a decrease of the number of α-syn molecules due to their transition to the retrograde state, and the third term characterizes an increase of the number of α-syn molecules due to their transition to the anterograde state (Fig. 2a).

Stating the conservation of α-syn in the retrograde state, characterized by concentration 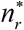 (Fig. 2a), we obtained the following equation:

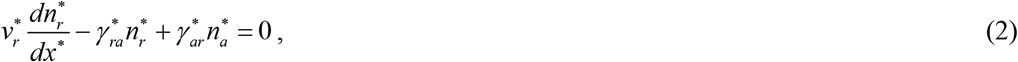

where 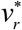 is the velocity of rapid motions of α-syn on MTs propelled by dynein motors in slow axonal transport. The first term on the left-hand side of Eq. (2) characterizes the effect of retrograde transport of α-syn due to the action of dynein motors while the second and third terms on the left-hand side are the kinetic terms characterizing the effects of α-syn transitions to the anterograde and retrograde kinetic states, respectively.

Eqs. (1) and (2) must be solved subject to the following boundary conditions. We assumed that α-syn is synthesized in the soma at a constant rate. Since all of the synthesized α-syn must enter the axon, this results in the following boundary condition at the axon hillock:

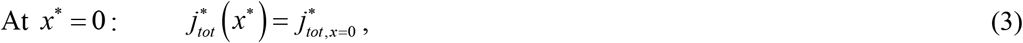

where 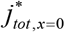 is the flux of α-syn at the axon hillock.

A given (high) concentration of α-syn, 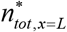, is imposed at the axon tip:

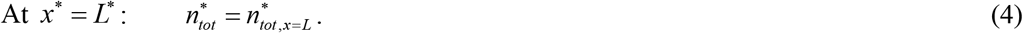

For the model displayed in Fig. 2a the total flux of cargo, 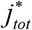, can be calculated as the difference between the anterograde and retrograde motor-driven fluxes:

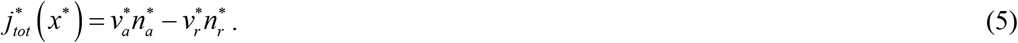

The total cargo concentration, 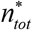, is found as the sum of cargo concentrations in the anterograde and retrograde kinetic states:

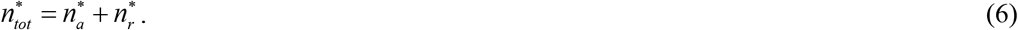

#### 2.1.2. A perturbation solution of the anterograde-retrograde cargo transport model for the case of small dynein velocity

The dimensionless retrograde velocity is defined as:

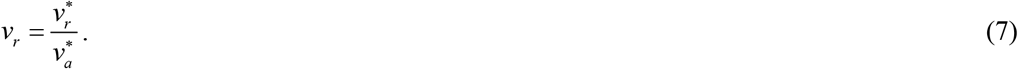

We are interested in investigating whether the model can simulate cargo transport against the cargo’s concentration gradient if dynein velocity is small compared to kinesin velocity. For this reason, we assumed that *v_r_* is a small parameter. The following perturbation expansions were utilized:

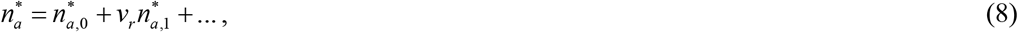

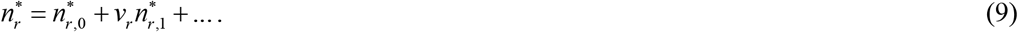

The substitution of Eqs. (8) and (9) into Eqs. (1) and (2) results in

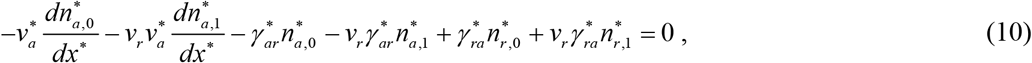

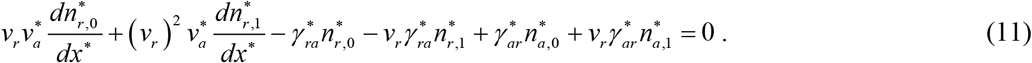

By separating the terms that do and do not contain the small parameter *v_r_* in Eqs. (10) and (11), and equating the terms that do not contain *v_r_* to zero, the following is obtained:

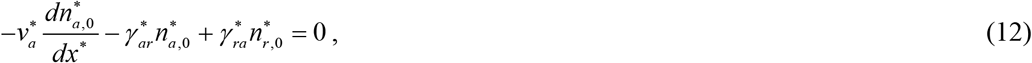

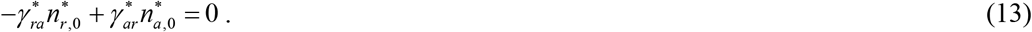

Solving Eq. (13) for 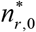, we obtained:

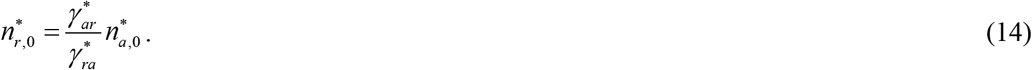

Eliminating 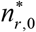 from Eq. (12), the following was obtained:

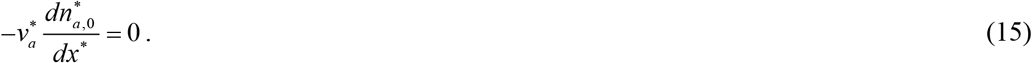

The solution of Eq. (15) is

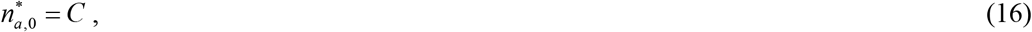

where *C* is the integration constant.

Substituting Eq. (16) into Eq. (14), we obtained that

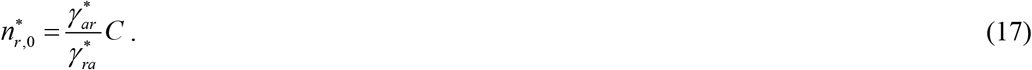

The utilization of Eqs. (16) and (17) gives the following equation for the total cargo concentration:

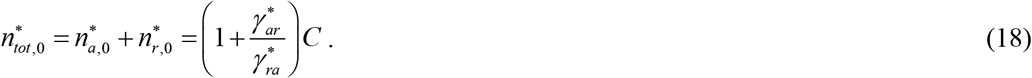

Using Eq. (3) and solving Eq. (18) for *C*, the following was obtained:

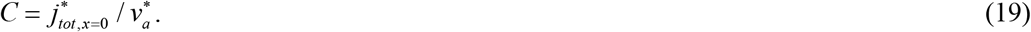

Substituting Eq. (19) into Eq. (16) gives that

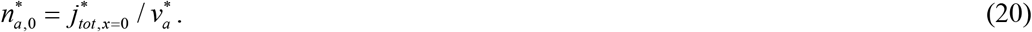

Substituting Eq. (19) into Eq. (17) results in the following:

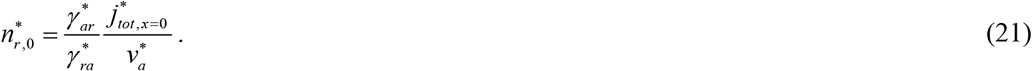

Finally, substituting Eq. (19) into Eq. (18) gives the following equation for the total cargo concentration:

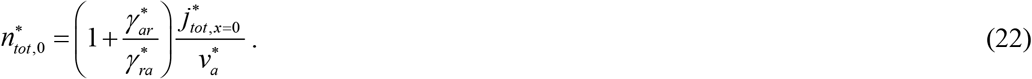

Eq. (22) shows that dynein dysfunction prevents the maintenance of a higher concentration of α-syn at the axon tip than at the soma.

### 2.2. Full slow axonal transport model

#### 2.2.1. General formulation of the full slow axonal transport model

We now move on to the full slow axonal transport model (kinetic diagram is shown in Fig. 2b). The continuum slow axonal transport model for simulating neurofilament transport was previously developed in [26]. The model was extended in [27] and [28] to simulate slow axonal transport of cytosolic proteins by adding a kinetic state for proteins that freely diffuse in the cytosol as well as a degradation term for the destruction of freely moving cytosolic proteins in proteasomes.

Conservation of α-syn driven anterogradely by kinesin motors is stated by the following equation:

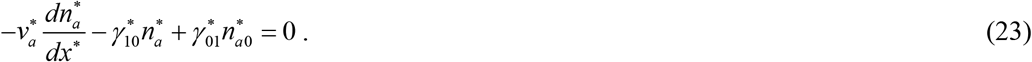

Eq. (23) is similar to Eq. (1), but kinetic constants are different and are now defined in Fig. 2b. The first term on the left-hand side of Eq. (23) describes the effect of kinesin-driven transport of α-syn while the second and third terms describe transitions between various kinetic states.

Conservation of α-syn driven retrogradely by dynein motors is stated by the following equation:

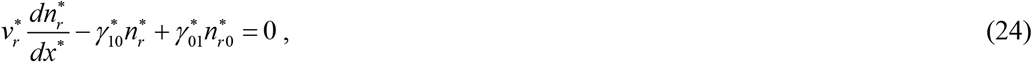

where the first term describes the effect of dynein-driven transport and the second and third terms are kinetic terms describing α-syn transitions between kinetic states. This equation is identical to Eq. (2), but kinetic constants are different.

Since cargos spend the majority of time in the pausing state in slow axonal transport [10], the full model also includes equations stating α-syn conservation in the pausing states. It is assumed that despite pausing α-syn retains its association with the motors and is ready to resume its motion in the anterograde or retrograde direction when the motors reconnect to MTs. The corresponding α-syn concentrations in the pausing states are 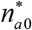 and 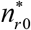, respectively (Fig. 2b):

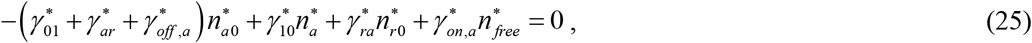

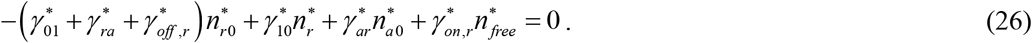

Because α-syn does not move in the pausing states, Eqs. (25) and (26) include only kinetic terms, which describe transitions between different kinetic states (Fig. 2b).

Stating conservation of α-syn in the free (cytosolic) state gives the following equation:

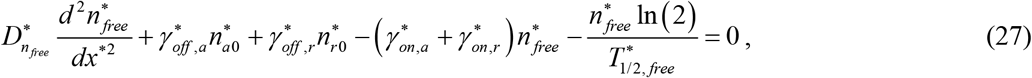

where 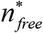 is the concentration of α-syn in the free (cytosolic) state, 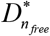 is the diffusivity of free α-syn, and 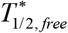 is the half-life of free α-syn. The first term on the left-hand side of Eq. (27) accounts for α-syn diffusion when α-syn is not connected to MTs. The presence of a diffusible fraction is typical for cytosolic proteins transported in SCb [29]. The last term on the left-hand side of Eq. (27) simulates α-syn destruction in proteasomes [30].

Note that in Eqs. (23)–(27) the degradation term is present only in the free state. Indeed, to enter a proteasome, α-syn must be detached from MTs. Thus, proteins moving on MTs are not subject to degradation in proteasomes, unlike those that are freely diffusing in the cytosol. This must be the case because in long neurons the time that it takes for a protein to travel from the soma to the axon tip exceeds the lifetime of free proteins. Therefore, it is likely that during their transit in axons proteins are protected from degradation [31,32]. If proteins can only be destroyed in the free state, this would explain how this protection is accomplished. Another possibility is that since cytosolic proteins are transported in SCb in the form of large cargo structures (multicargo complexes), they may be too large to enter proteasomes and cannot be degraded at all during transport. In this case, the half-life of α-syn complexes in the free state should be close to infinity.

If α-syn diffusivity 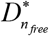 in Eq. (27) is not zero, Eqs. (23)–(27) must be solved subject to four boundary conditions. At the axon hillock, we imposed the total concentration of α-syn at the hillock, 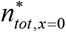, and the flux of α-syn entering the axon, which equals to 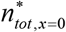 times the average velocity of α-syn (which includes pauses), 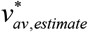.

At *x*\* =0:

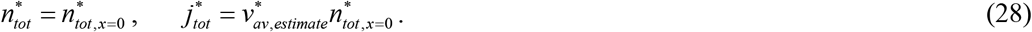

The total flux of α-syn in the full slow axonal transport model equals to the sum of diffusion-driven α-syn flux, the kinesin-driven anterograde flux, and the dynein-driven retrograde flux, respectively:

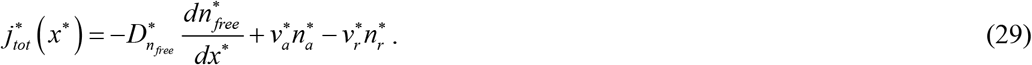

The total concentration of α-syn now equals the sum of α-syn concentrations in all five kinetic states displayed in Fig. 2b:

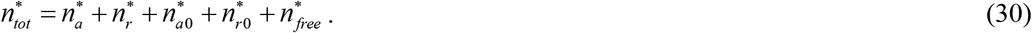

The average velocity of α-syn (a parameter that depends on *x*\*, see ref. [33]) can then be calculated as:

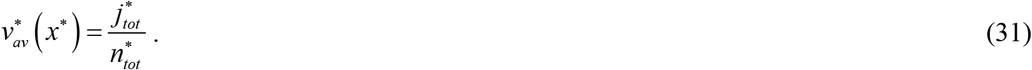

At the axon tip, we imposed a zero gradient of the free α-syn concentration (which is equivalent to a zero α-syn diffusion flux) and a given total α-syn concentration, 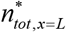.

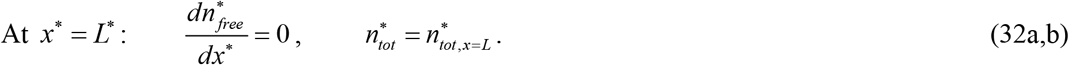

It is convenient to define the dimensionless concentration (for all α-syn concentration components) as:

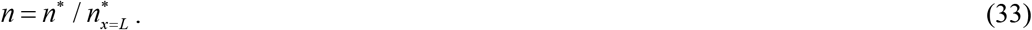

This allows recasting Eq. (32b) as follows:

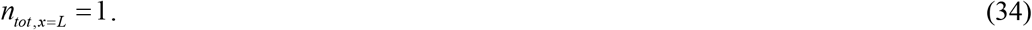

The dimensionless total flux of α-syn is then defined as follows:

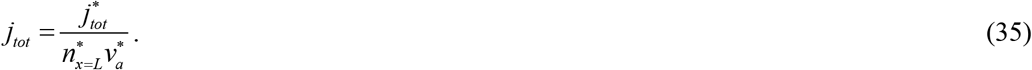

#### 2.2.2. A perturbation solution of the full slow axonal transport model for the case of small cargo diffusivity and small dynein velocity

Unless proteins are MT-bound, such as tau or MAP2, slow axonal transport cargos are usually not transported along the axon as single particles but in protein complexes [10,13,29,34–38]. Because diffusivity of such multiprotein complexes is expected to be very small, we investigated how Eqs. (23)–(27) can be simplified for the limiting case of vanishing diffusivity of free proteins. Our goal is to understand whether after such simplification the model is capable of simulating cargo transport against a cargo concentration gradient.

The dimensionless cargo diffusivity is defined as:

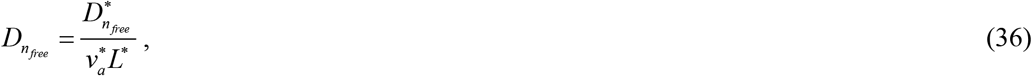

which is reciprocal to the Peclet number. Since *D_n_free__* can be re-written as:

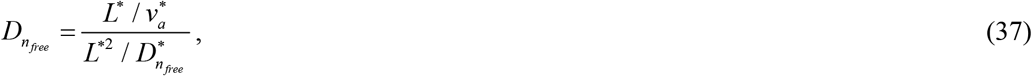

it can be viewed as the ratio of the convection time to diffusion time.

We assumed that *D_n_free__* is a small parameter. The following perturbation expansions are utilized:

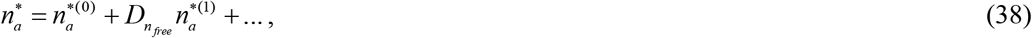

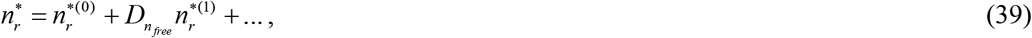

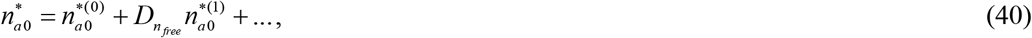

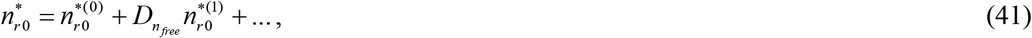

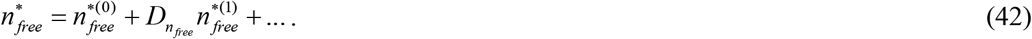

Using Eqs. (40)–(42), Eq. (27) is recast as:

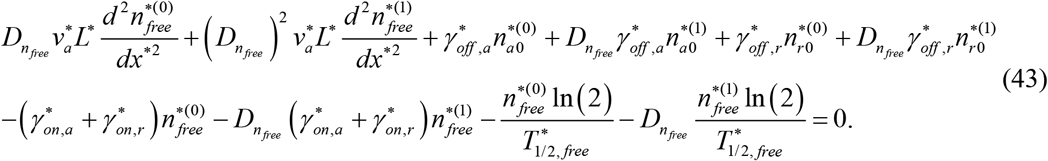

By separating the terms that do and do not contain the small parameter *D_n_free__*, and equating the terms that do not contain *D_n_free__* to zero, the following is obtained:

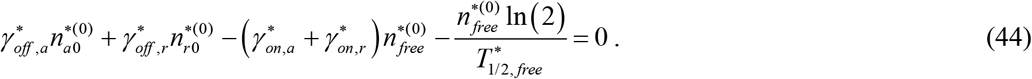

By using Eq. (38)–(42) in Eq. (23)–(26), separating the terms that do and do not contain the small parameter *D_n_free__*, and equating the terms that do not contain *D_n_free__* to zero, the following is obtained:

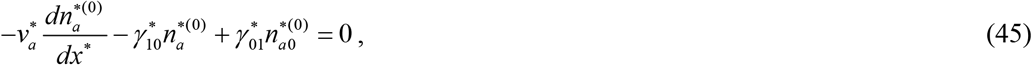

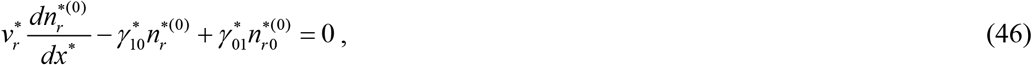

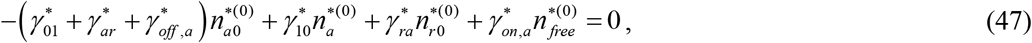

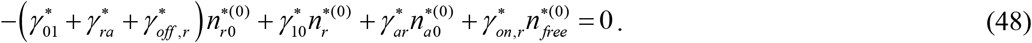

Because we investigate the case of dynein dysfunction, which we simulate by a vanishing dynein velocity, we also assumed that *v_r_* defined in Eq. (7) is a small parameter. The following perturbation expansions are utilized:

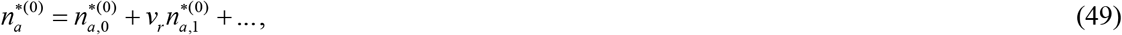

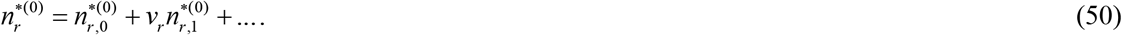

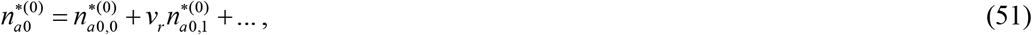

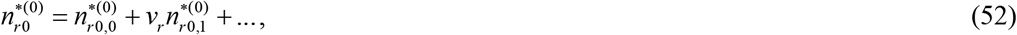

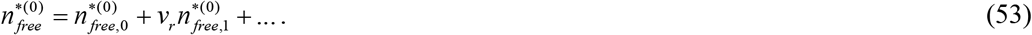

By using Eqs. (49)–(53) in Eqs. (44)–(48), separating the terms that do and do not contain the small parameter *v_r_*, and equating the terms that do not contain *v_r_* to zero, the following is obtained:

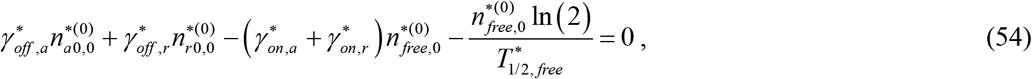

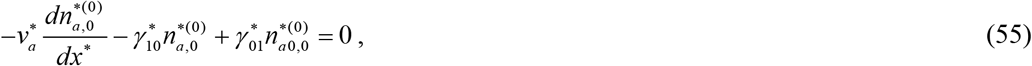

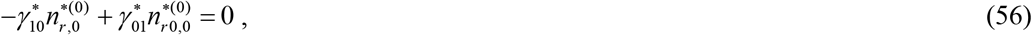

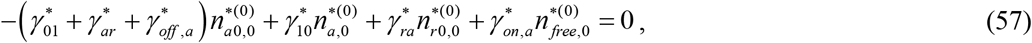

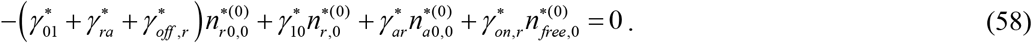

From Eq. (56):

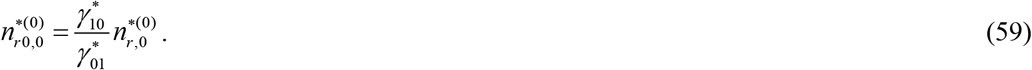

Eqs. (54), (57), and (58) are then recast as:

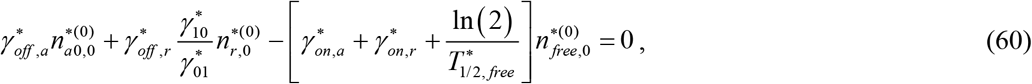

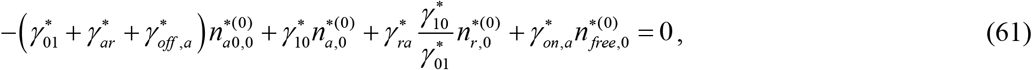

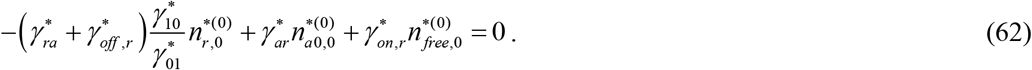

Eliminating 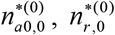, and 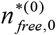 from Eqs. (55), (60)–(62), the following equation for 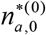 was obtained:

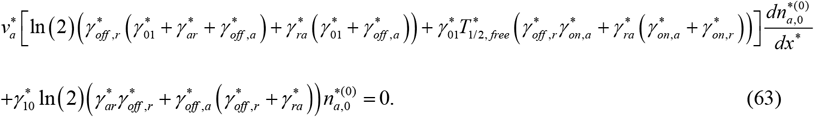

For the case when 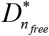 and 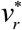 are small, the flux of cargos is given by

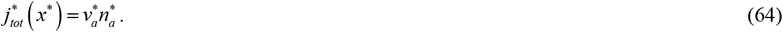

Eq. (63) is solved subject to the following boundary condition, which requires the flux of α-syn entering the axon to be the same as in the full slow axonal transport model:

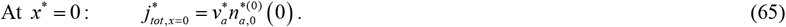

The solution of Eq. (63) with boundary condition (65) is

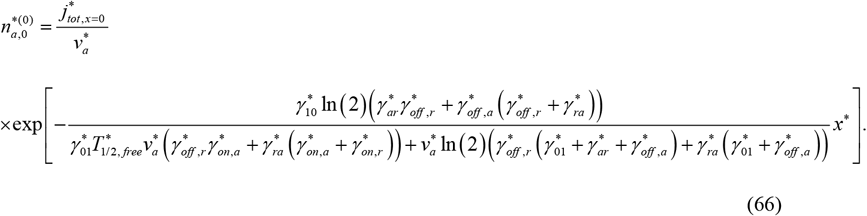

Note that the solution given by Eq. (66) has the form 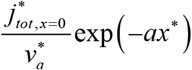, where the concentration decay constant *a* > 0, which means that 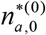 cannot describe an increasing concentration toward the terminal.

If *a* is small, the solution is 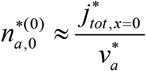, which is a constant value.

By eliminating 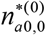 and 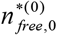 from Eqs. (60)–(62), the following equation for 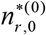 is obtained:

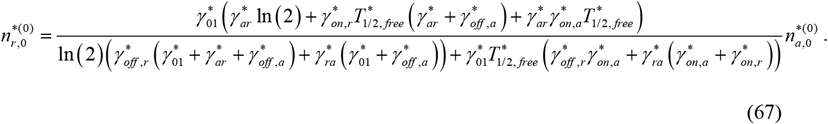

By eliminating 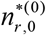 and 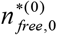 from Eqs. (60)–(62), the following equation for 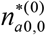 is obtained:

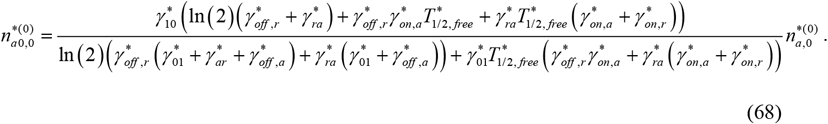

By eliminating 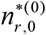 and 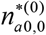 from Eqs. (60)–(62), the following equation for 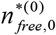 is obtained:

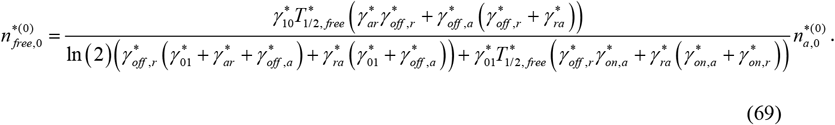

The total concentration, 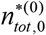, can be obtained by finding the sum of concentration components given by Eqs. (59) and (66)–(69).

### 2.3. Sensitivity of the concentration boundary layer thickness to dynein velocity

The thickness of the concentration boundary layer is defined as the distance from the axon tip to the location 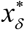 (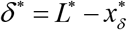, see Fig. 4) where the total cargo concentration drops by 99%, which means where it reaches the value

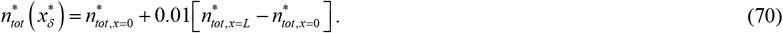

The sensitivity of *δ*\* to the dynein velocity 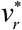 is calculated as follows [39–42]:

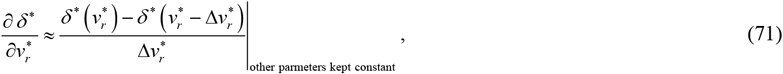

where 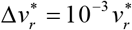 is the step size. The independence of the sensitivity to the step size was tested by varying the step sizes.

**Fig. 3.**
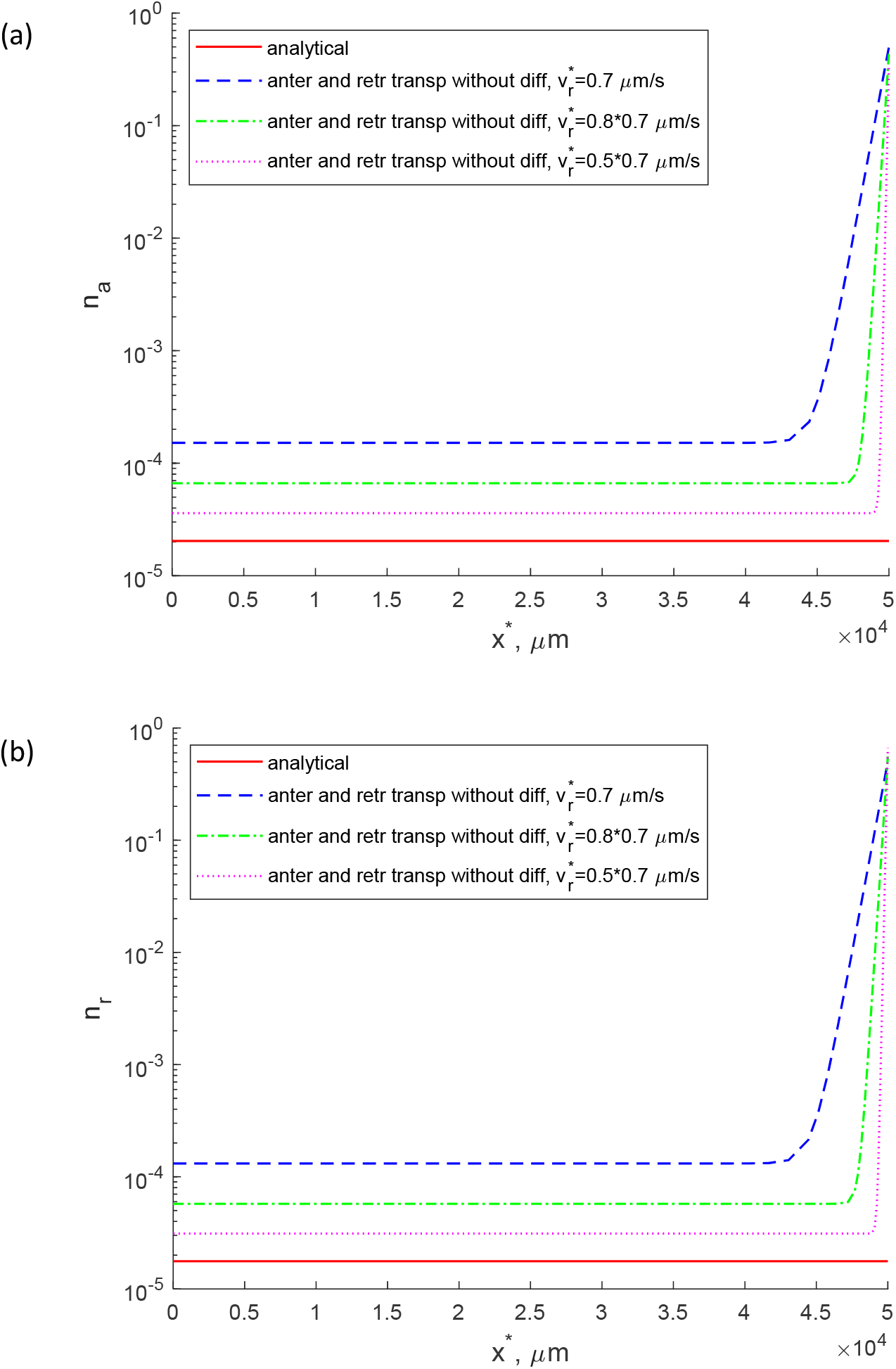
Anterograde and retrograde axonal transport model without pausing states and diffusion (kinetic diagram for this model is shown in Fig. 2a). (a) Concentration of cargos transported by anterograde motors. (b) Concentration of cargos transported by retrograde motors.

**Fig. 4.**
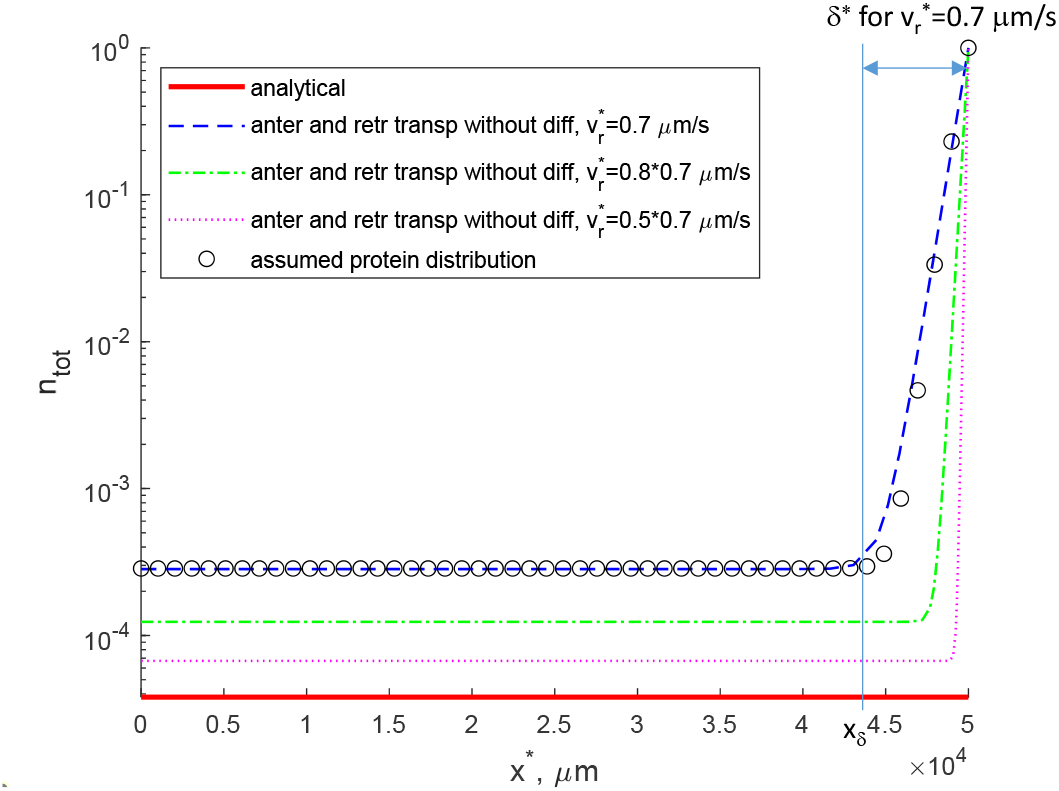
Anterograde and retrograde axonal transport model without pausing states and diffusion (kinetic diagram for this model is shown in Fig. 2a). Total cargo concentration (the sum of cargo concentrations in anterograde and retrograde motor-driven states).

To make the sensitivity independent of parameter magnitude, the non-dimensional relative sensitivity was calculated as [40,43]:

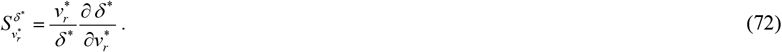

### 2.4. Finding best-fit values of kinetic constants by multi-objective optimization

To solve the anterograde-retrograde transport problem given by Eqs. (1) and (2) with boundary conditions (3) and (4) values of kinetic constants 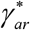 and 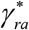 are needed. To solve the full slow axonal transport problem given by Eqs. (23)–(27) with boundary conditions (28) and (32), values of all eight kinetic constants given in Table S3 must be determined. We determined best-fit values of these kinetics constants by multi-objective optimization [44–48]. We minimized the following objective (penalty) function, which combines three different effects:

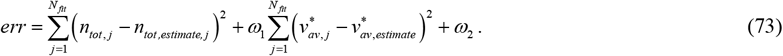

The first term on the right-hand side of Eq. (73) minimizes the difference between the computed concentration, *n_tot, j_*, and the synthetic data, *n_tot, estimate, j_*, in *N_fit_* uniformly spaced points along the length of the axon. We know that α-syn predominantly localizes in the presynaptic terminal [49–51]. To simulate this, we assumed that the concentration of synthetic data is given by the following modified logistic function:

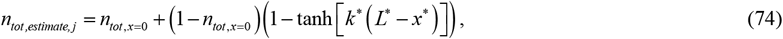

where *k*\* is the logistic growth rate, a parameter that characterizes the steepness of the *S*-shaped curve used in approximating the distribution of α-syn along the axon length. We used *k*\* = 10^-3^ 1/μm. The distribution given by Eq. (74) is shown by hollow circles in Fig. 4.

The purpose of the second term on the right-hand side of Eq. (73) is to simulate the difference between the numerically predicted α-syn velocity, 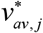, and experimentally reported α-syn velocity, 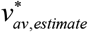. We used 0.05 μm/s for 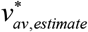, a value which is in the middle of the experimentally reported range of SCb velocity, 2-8 mm/day [13]. In the second term, we also used *ω*_1_=1 s^2^/μm^2^. This value was selected based on numerical experimentation, to avoid overfitting either α-syn concentration or its average velocity, see ref. [52].

The third term on the right-hand side of Eq. (73) is set to a large value, *ω*_2_ = 10^8^, if any of α-syn concentration components, *n_a, j_*, *n_r, j_*, *n*_*a*0,*j*_, *n*_*r*0,j_, *n_free, j_*, or α-syn flux, *j_tot_* (*j*=1,…,*N_fit_*) becomes negative. The latter condition comes from the assumption that α-syn is not synthesized at the terminal, and that the half-life of α-syn is finite; therefore, the terminal must be supplied by α-syn from the soma. Best-fit values of kinetic constants for the anterograde-retrograde transport model are given in Table S4, and the best-fit values of kinetic constants for the full slow axonal transport model are given in Table S5.

### 2.5. Numerical solution

Eqs. (1) and (2) were solved using MATLAB’s BVP5C solver (MATLAB R2020b, MathWorks, Natick, MA, USA). For the full slow axonal transport model, we first eliminated 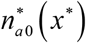 and 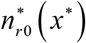 from Eqs. (23)–(27) using Eqs. (25) and (26). The resulting system of ordinary differential equations of the fourth order was again solved using MATLAB’s BVP5C solver. To determine values of kinetic constants (Tables S4 and S5), we used MULTISTART with the local solver FMINCON. These routines are part of MATLAB’s Optimization Toolbox. We used 1000 random points plus the initial point given in Table S3 as starting points in searching for the global minimum. We followed the numerical procedure described in ref. [53].

## 3. Results

### 3.1. Anterograde and retrograde axonal transport model without pausing states and diffusion fails to simulate an increase of cargo concentration toward the axon tip if dynein velocity is small

We used the anterograde-retrograde model (Fig. 2a) to simulate transport of an SCb protein that is predominantly localized in the presynaptic terminal. α-syn data were used for our simulation. We found that the concentrations of proteins pulled by anterograde and retrograde motors remain constant along most of the axon length and then sharply increase near the axon tip (Fig. 3a,b). A decrease of the dynein velocity results in a decrease of the boundary layer thickness (i.e. the region where cargo concentrations change from the constant low concentration which is maintained along most of the axon length to the high concentration seen at the axon tip; Fig. 3a,b). If the dynein velocity approaches zero, the anterograde and retrograde concentrations are represented by horizontal lines (no concentration increase toward the axonal tip, see the curves corresponding to the analytical solution in Fig. 3a,b).

A similar decrease of the boundary layer thickness with a decrease of the dynein velocity is observed in Fig. 4 which displays the total cargo concentration. In the model given by Eqs. (1) and (2) the boundary layer thickness is very sensitive to the dynein velocity. The dimensionless sensitivity, 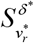, calculated using Eq. (72) for this model is 9.21. Therefore, in addition to the base case, corresponding to 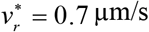, we show two more concentration distributions where the dynein velocity is decreased by factors of 0.8 and 0.5 (see Figs. 3 and 4). Notably, further decreasing dynein velocity causes the boundary layer thickness to become small, leading to an unphysiological near-vertical transition in concentration distribution.

The values of kinetic constants used for computing all curves in Figs. 3, 4, and S1 were determined using 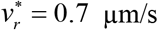. Otherwise, if kinetic constants were optimized for each value of 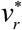, all three numerical curves would be the same and would give a perfect fit with the synthetic data. Numerically computed curves in Figs. 3 and 4 exhibit a constant concentration along most of the axonal length which is followed by a sharp concentration increase near the axon tip. This feature is due to the concentration distribution of synthetic data (see Eq. (74)). The sharp increase of the total concentration near the terminal exhibited by synthetic data (see hollow circles in Fig. 4) simulates the presynaptic localization of α-syn. Since values of kinetic constants in the model are determined such that numerically predicted total cargo concentration would fit the synthetic data distribution, the shapes of the numerical curves in Fig. 4 correspond to the synthetic data distribution. The important feature in Fig. 4 is the inability of the model to simulate the increase of cargo concentration toward the axonal tip if dynein velocity tends to zero (see the curve that represents the analytical solution).

The total flux of cargos (defined in Eq. (5)) is constant independent of the position in the axon (Fig. S1) because the cargo flux at the hillock is imposed by Eq. (3) and there is no cargo degradation in the model given by Eqs. (1) and (2).

### 3.2. For small cargo diffusivity, the full slow axonal transport model fails to simulate an increase of cargo concentration toward the axon tip if dynein velocity is small

For the full slow axonal transport model (Eqs. (23)–(27), Fig. 2b) we observed that predicted cargo concentrations are less sensitive to the dynein velocity than in the model given by Eqs. (1) and (2). The dimensionless sensitivity, 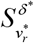, calculated using Eq. (72) for the full slow axonal transport model is 1.28. Therefore, in Figs. 5, 6, and S2–S4 we have shown results for the base dynein velocity 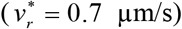 and two much smaller values of dynein velocity, 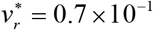 and 0.7 × 10 ^2^ μm/s. Again, the values of kinetic constants used for computing all curves in Figs. 5, 6, and S2–S4 were determined using 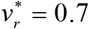. The analytical solution obtained for the case when the cargo diffusivity and dynein velocity both approach zero predicts almost uniform cargo concentrations (Figs. 5, 6, S2, and S3a). Indeed, for the results displayed in Figs. 5 and 6 the concentration decay constant *a* defined in the paragraph after Eq. (66) equals to 3.59 × 10^-13^ 1/μm. This means that the exponential function in Eq. (66) is almost identical to unity and 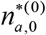 is constant. This explains why the model predicts that without a retrograde motor neurons are incapable of supporting high cargo concentration at the axon terminal. It should be noted that even if the soma can sense the low cargo concentration at the axon terminal (if some retrograde signaling is present) and starts producing more α-syn, it will only shift the horizontal “analytical* α-syn distribution displayed in Fig. 6 upward, but the distribution will remain horizontal.

**Fig. 5.**
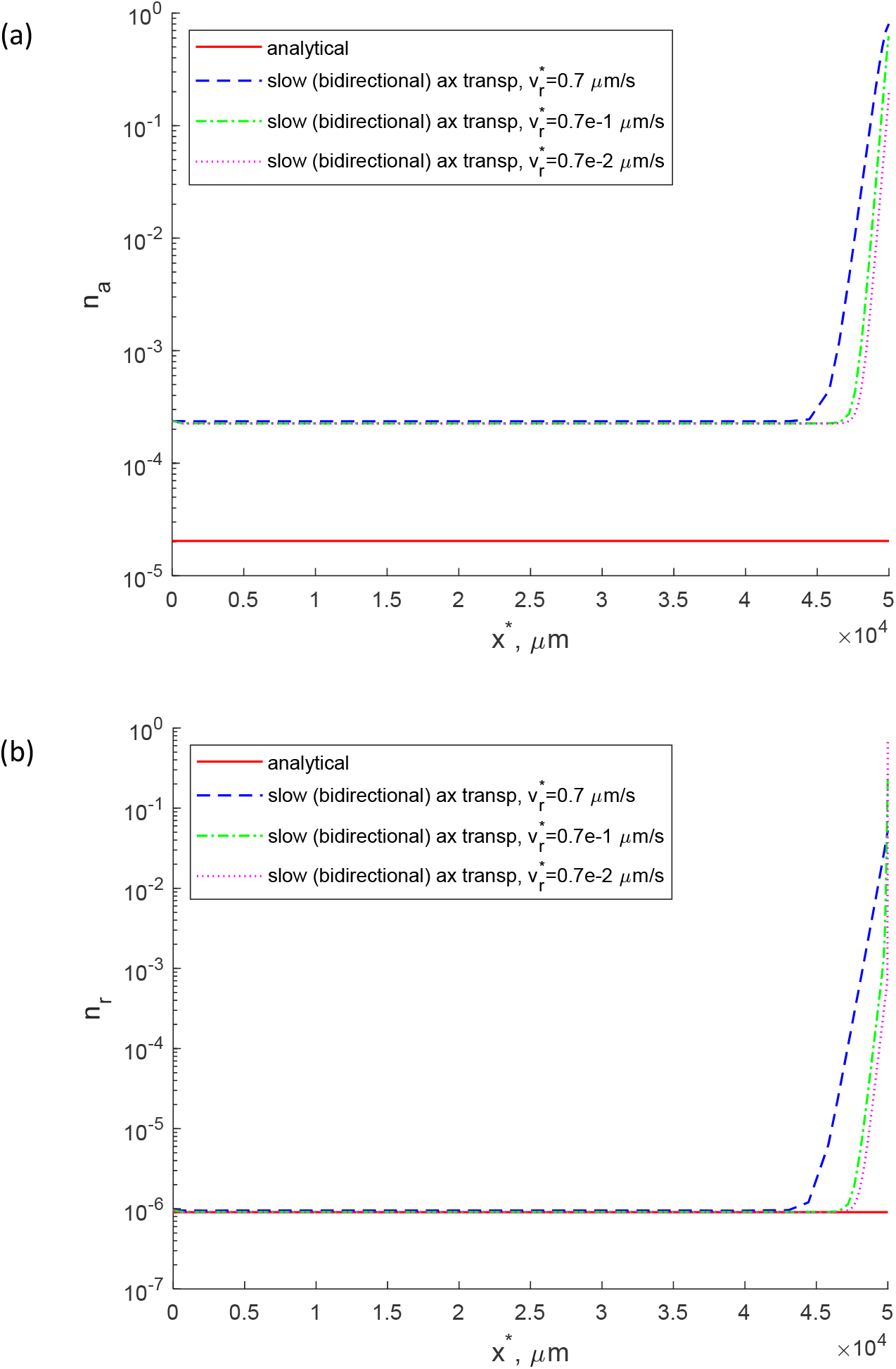
Full slow axonal transport model (kinetic diagram for this model is shown in Fig. 2b). (a) Concentration of cargos transported by anterograde motors. (b) Concentration of cargos transported by retrograde motors. 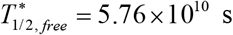.

**Fig. 6.**
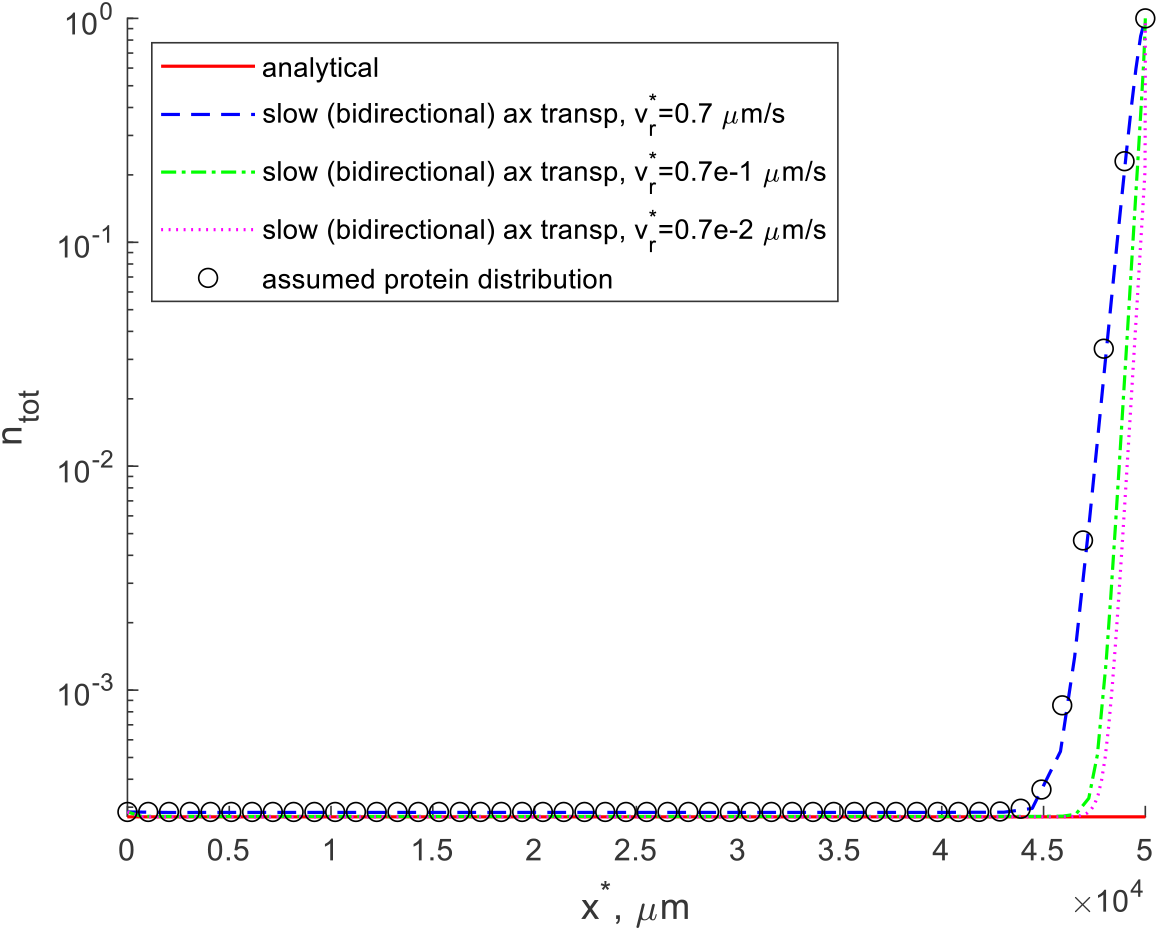
Full slow axonal transport model (kinetic diagram for this model is shown in Fig. 2b). Total cargo concentration (the sum of cargo concentrations in motor-driven, pausing, and diffusion-driven states). 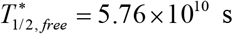.

For the full slow axonal transport model, the cargo flux displayed in Fig. S3b remains constant because of the large value of cargo half-life used in computations (Fig. S3b is computed for 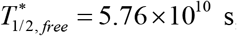, assuming that α-syn is protected from degradation during its transport in the axon). Computations using a smaller cargo half-life 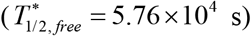 predict a decay of the total flux of cargos near the axon tip (Fig. S4b). This is because the model assumes that α-syn can be destroyed in proteasomes only in the free (cytosolic) state. Since the concentration of free α-syn increases toward the tip (Fig. S4a), there is more α-syn degradation near the tip. Hence, the total flux of α-syn decreases near the tip (Fig. S4b).

## 4. Discussion, limitations of the model, and future directions

The results obtained in this paper suggest that dynein dysfunction results in the inability of neurons to transport cargo against a concentration gradient. This result holds when cargo diffusivity is small, which applies to cargos transported as polymers or as a part of multiprotein complexes. A mathematical argument to explain why dynein motors are needed to support a cargo concentration distribution that increases toward the axon tip is as follows. With the anterograde-only motion of cargo there is no way to impose a boundary condition at the axon tip that is to require an elevated cargo concentration at the tip. Our results, counter-intuitively, suggest that the progressive distal to proximal (dying-back) axonal degeneration may result from dynein dysfunction because of the inability of the neuron to support a high cargo concentration at the terminal.

Limitations of our model are related to neglecting local protein synthesis and local controlled protein degradation. These effects should be included in future models and considered in future research.

Our results are testable and falsifiable. Future work could test whether neurons with dysfunctional dynein are indeed unable to support a cargo distribution in an axon that increases with distance from the soma. Ref. [11] reported that dynein knockdown causes arrest of bidirectional peroxisome transport in *Drosophila*. However, cargo transport can be rescued by artificially attaching a construct containing another retrograde motor (kinesin-14, Ncd) to peroxisomes. It would be interesting to investigate whether dysfunctional dynein can be replaced by another retrograde motor to support slow axonal transport.

## Acknowledgment

IAK acknowledges the fellowship support of the Paul and Daisy Soros Fellowship for New Americans and the NIH/National Institute of Mental Health (NIMH) Ruth L. Kirchstein NRSA (F30 MH122076-01). AVK acknowledges the support of the National Science Foundation (award CBET-2042834) and the Alexander von Humboldt Foundation through the Humboldt Research Award.

## Supplemental Materials

### S1. Supplementary tables

**Table S1.** Dependent variables in the full slow axonal transport model of α-syn transport from the soma to the axon tip (Fig. 2b). The model of anterograde-retrograde transport displayed in Fig. 2a contains only two concentrations, 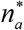 and 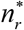.

| Symbol | Definition | Units |
| --- | --- | --- |
| $n_a^*$ | Concentration of on-track α-syn moving along MTs anterogradely, propelled by molecular motors | $\mu\text{m}^{-1}$ |
| $n_r^*$ | Concentration of on-track α-syn moving along MTs retrogradely, propelled by molecular motors | $\mu\text{m}^{-1}$ |
| $n_{a0}^*$ | Concentration of pausing on-track α-syn monomers that are still associated with molecular motors and can resume their anterograde motion | $\mu\text{m}^{-1}$ |
| $n_{r0}^*$ | Concentration of pausing on-track α-syn monomers that are still associated with molecular motors and can resume their retrograde motion | $\mu\text{m}^{-1}$ |
| $n_{free}^*$ | Concentration of free (off-track) α-syn in the cytosol | $\mu\text{m}^{-1}$ |

**Table S2.** Parameters characterizing transport of α-syn in the axon obtained from published data or assumed on physical grounds.

| Symbol | Definition | Units | Value or range | Reference(s) | Value(s) used in computations |
| --- | --- | --- | --- | --- | --- |
| $D_{n_{free}}^*$ | Diffusivity of monomeric α-syn in the free (cytosolic) state | $\mu\text{m}^2/\text{s}$ | 114 <sup>a</sup> | [54] | 0, 114 |
| $k^*$ | The logistic growth rate, which characterizes the steepness of the $S$-shaped curve used in approximating the distribution of α-syn along the axon length | $1/\mu\text{m}$ | | | $10^{-3}$ |
| $L^*$ | Length of the axon | $\mu\text{m}$ | up to $10^6$ in humans | [38] | $5 \times 10^4$ <sup>b</sup> |
| $n_{tot,x=0}^*$ | Concentration of α-syn at the axon hillock | $\mu\text{M}$ | $10^{-3}$ <sup>c</sup> | [55] | $10^{-3}$ |
| $n_{tot,x=L}^*$ | Concentration of α-syn at the presynaptic terminal | $\mu\text{M}$ | 3.5 <sup>c</sup> | [56] | 3.5 |
| $N_{fit}$ | Number of points uniformly distributed along the axon length in which we matched the α-syn concentration with the assumed $S$-shaped distribution | | | | 50 |
| $T_{1/2,free}^*$ | Half-life of free α-syn | s | $6.62 \times 10^3$ - $1.73 \times 10^5$ <sup>d</sup> | [57-60] | $5.76 \times 10^4$, $5.76 \times 10^{10}$ |
| $v_a^*$ | Velocity of rapid motions of α-syn on MTs propelled by kinesin motors in slow axonal transport | $\mu\text{m s}^{-1}$ | 0.7 | [61] | 0.7 |
| $v_r^*$ | Velocity of rapid motions of α-syn on MTs propelled by dynein motors in slow axonal transport | $\mu\text{m s}^{-1}$ | 0.7 | [61] | 0, $0.7 \times 10^{-2}$, $0.7 \times 10^{-1}$, $0.5 \times 0.7$, $0.8 \times 0.7$, 0.7 |
| $v_{av,estimate}^*$ | Estimated average velocity of α-syn in slow axonal transport (assumed to be constant) | $\mu\text{m s}^{-1}$ | 0.023-0.093 <sup>e</sup> | [13,31,62] | 0.05 |
| $\omega_1$ | Weighting factor in Eq. (73) | $\text{s}^2/\mu\text{m}^2$ | | Numerical experimentation | 1 |
| $\omega_2$ | Weighting factor in Eq. (73) | | | Numerical experimentation | $10^8$ |

<sup>a</sup> Monomeric α-syn has large diffusivity, which is explained by its small molecular weight of $\sim 14$ kDa [63].
<sup>b</sup> We used a representative axon length of 50 mm [38].
<sup>c</sup> According to [55], the α-syn concentration in presynaptic terminals is estimated to be in the micromolar range. In other parts of the cell, the α-syn concentration can be as low as in the nanomolar range [55]. According to [56], the concentration of α-syn in the presynaptic terminal is estimated to be $3.5 \mu\text{M}$ .
<sup>d</sup> Cytosolic proteins are usually degraded in proteasomes. Their typical half-life is days or weeks at the most [64].
It is likely that monomeric α-syn is degraded in proteasomes [57,65]. Estimates reported in the literature include 1.84 hours [57], 16 hours [58], and 48 hours [59]. We followed [58,60], and used 16 hours ( $5.76 \times 10^4$ s) as an estimate of the half-life of an unprotected α-syn.
In long neurons, the time that it takes for a protein to travel from the soma to the synapse may exceed the lifetime of the monomeric protein. Therefore, it is likely that proteins are somehow protected from degradation during their transit in the axon [31,32]. As a result, the half-life of α-syn undergoing slow axonal transport may be much larger than 16 hours. This could be explained by transport in large multiprotein complexes [10] which are too large to enter a proteasome for degradation. For this reason, in computations we used two estimates for α-syn half-life: $5.76 \times 10^4$ s (16 hours) and $5.76 \times 10^{10}$ s ( $1.6 \times 10^7$ hours).
<sup>e</sup> Several different estimates of the average velocity of α-syn transport in axons have been reported: $0.022$ - $0.028 \mu\text{m/s}$ ( $1.9$ - $2.4$ mm/day) [62] and $0.035$ - $0.12 \mu\text{m/s}$ ( $3$ - $10$ mm/day) [31]. Since α-syn is transported in SCb, we used an estimate reported in [13], $0.023$ - $0.093 \mu\text{m/s}$ ( $2$ - $8$ mm/day).

**Table S3.** Kinetic constants characterizing transition of α-syn between different kinetic states in the slow axonal transport model, which are displayed in Fig. 2b. The anterograde-retrograde model displayed in Fig. 2a includes only two kinetic constants, 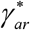 and 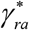. The values of these kinetic constants were determined by using multi-objective optimization to find values that give the best fit with synthetic data, as described in section 2.4, see also [39,52,66].

| Symbol | Definition | Units | Initial guess used in the optimization procedure |
| --- | --- | --- | --- |
| $\gamma_{10}^*$ | Kinetic constant describing the rate of transitions $n_a^* \rightarrow n_{a0}^*$ and $n_r^* \rightarrow n_{r0}^*$ (Fig. 2b) | $s^{-1}$ | $7.19 \times 10^{-3}$ |
| $\gamma_{01}^*$ | Kinetic constant describing the rate of transitions $n_{a0}^* \rightarrow n_a^*$ and $n_{r0}^* \rightarrow n_r^*$ (Fig. 2b) | $s^{-1}$ | $4.01 \times 10^{-3}$ |
| $\gamma_{ar}^*$ | Kinetic constant describing the rate of transitions $n_{a0}^* \rightarrow n_{r0}^*$ (Fig. 2b) and $n_a^* \rightarrow n_r^*$ (Fig. 2a) | $s^{-1}$ | $4.59 \times 10^{-3}$ |
| $\gamma_{ra}^*$ | Kinetic constant describing the rate of transitions $n_{r0}^* \rightarrow n_{a0}^*$ (Fig. 2b) and $n_r^* \rightarrow n_a^*$ (Fig. 2a) | $s^{-1}$ | $6.76 \times 10^{-3}$ |
| $\gamma_{on,a}^*$ | Kinetic constant describing the rate of transitions $n_{free}^* \rightarrow n_{a0}^*$ (Fig. 2b) | $s^{-1}$ | $9.43 \times 10^{-2}$ |
| $\gamma_{on,r}^*$ | Kinetic constant describing the rate of transitions $n_{free}^* \rightarrow n_{r0}^*$ (Fig. 2b) | $s^{-1}$ | $5.90 \times 10^{-2}$ |
| $\gamma_{off,a}^*$ | Kinetic constant describing the rate of transitions $n_{a0}^* \rightarrow n_{free}^*$ (Fig. 2b) | $s^{-1}$ | $5.29 \times 10^{-3}$ |
| $\gamma_{off,r}^*$ | Kinetic constant describing the rate of transitions $n_{r0}^* \rightarrow n_{free}^*$ (Fig. 2b) | $s^{-1}$ | $5.40 \times 10^{-3}$ |

**Table S4.**
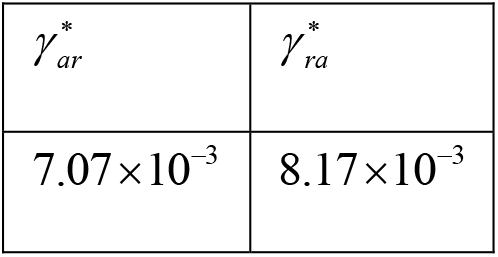
Kinetic constants characterizing the transition of α-syn between different kinetic states in an axonal transport model that includes anterograde and retrograde motor-driven kinetics states only (Fig. 2a). 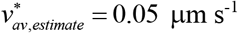. 1000 random points plus the initial point given in Table S3 (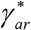 and 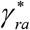 values only) were utilized as starting points in the search for the global minimum of the penalty function given by Eq. (73). All kinetic constants in Table S4 have dimensions s^-1^.

|  |  |
| --- | --- |
| $\gamma_{ar}^*$ | $\gamma_{ra}^*$ |
| $7.07 \times 10^{-3}$ | $8.17 \times 10^{-3}$ |

**Table S5.** Kinetic constants characterizing the transition of α-syn between different kinetic states in the full slow axonal transport model (Fig. 2b). 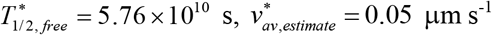, and 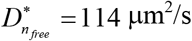. 1000 random points plus the initial point given in Table S3 were utilized as starting points in the search for the global minimum of the penalty function given by Eq. (73). All kinetic constants in Table S5 have dimensions s^-1^.

|  |  |  |  |  |  |  |  |
| --- | --- | --- | --- | --- | --- | --- | --- |
| $\gamma_{10}^*$ | $\gamma_{01}^*$ | $\gamma_{ar}^*$ | $\gamma_{ra}^*$ | $\gamma_{on,a}^*$ | $\gamma_{on,r}^*$ | $\gamma_{off,a}^*$ | $\gamma_{off,r}^*$ |
| $1.82 \times 10^{-3}$ | $1.65 \times 10^{-4}$ | $1.44 \times 10^{-4}$ | $5.87 \times 10^{-3}$ | $1.41 \times 10^{-1}$ | $2.49 \times 10^{-2}$ | $9.67 \times 10^{-3}$ | $3.53 \times 10^{-2}$ |

## S2. Supplementary figures

### S2.1. Supplementary figures for the simplified anterograde-retrograde axonal transport model

**Fig. S1.**
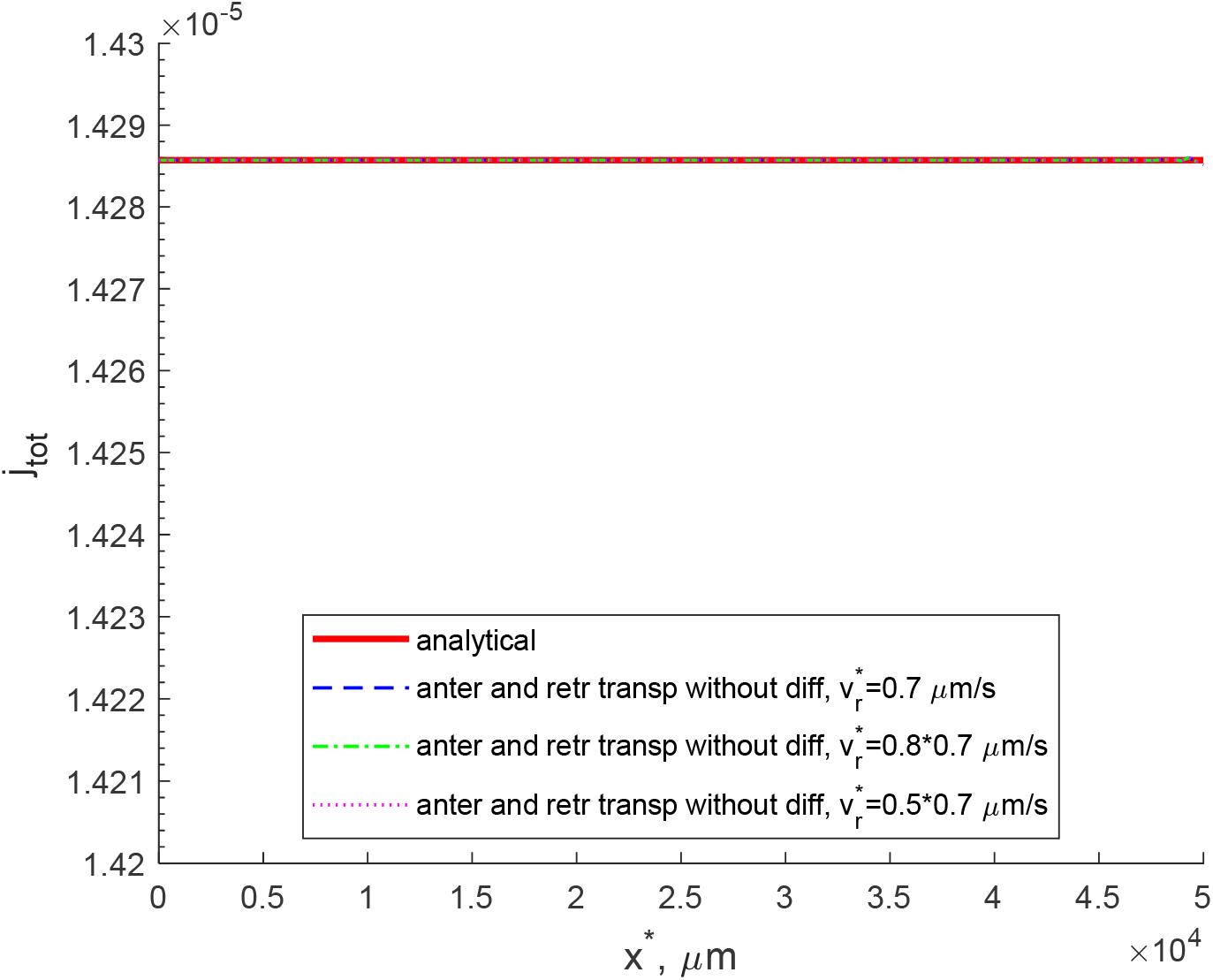
Anterograde and retrograde axonal transport model without pausing states and diffusion. Total flux of cargos due to the action of anterograde and retrograde motors. Note that the model does not include destruction of cargos in proteasomes, which explains the uniform flux of cargos.

### S2.2. Supplementary figures for the full slow axonal transport model

**Fig. S2.**
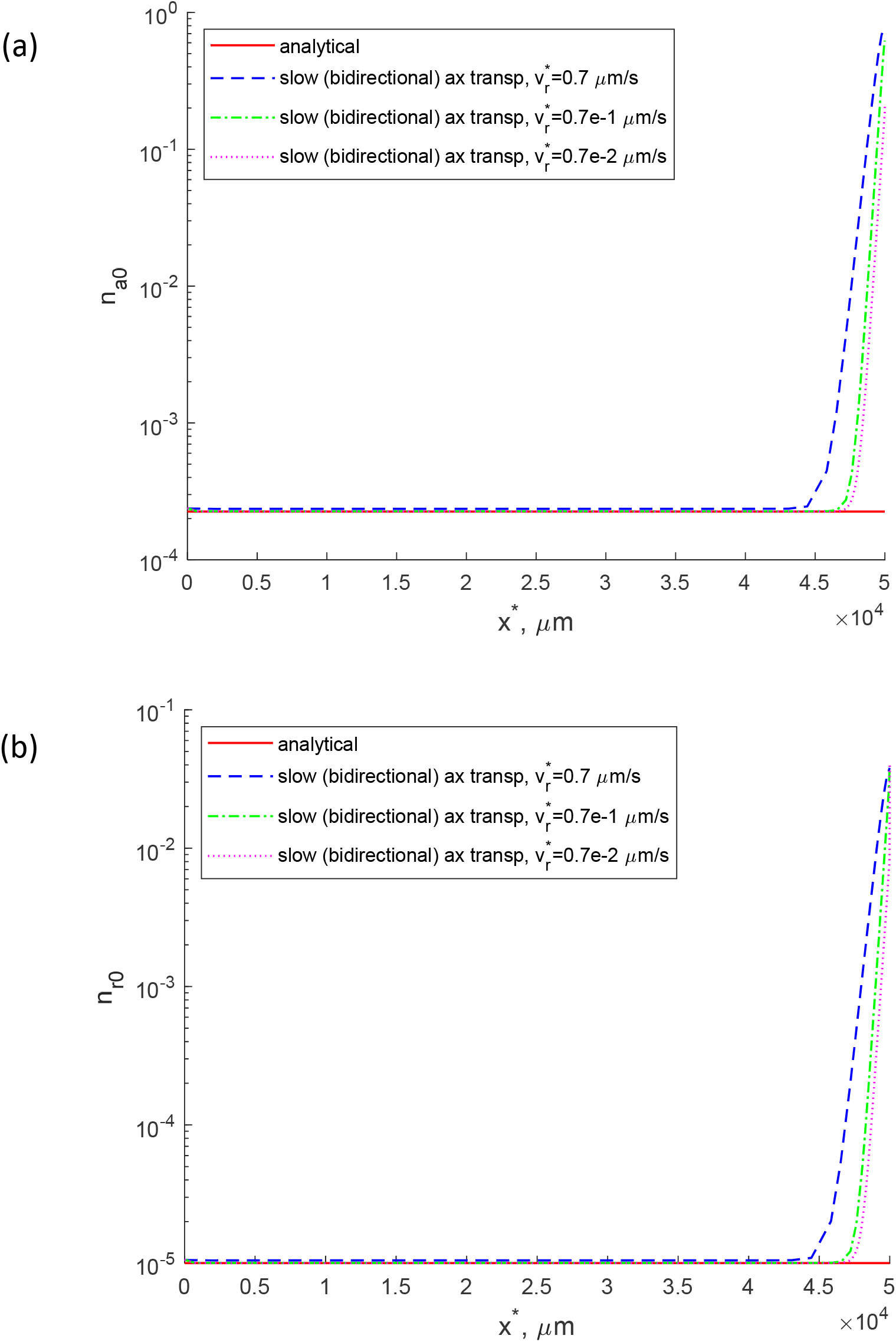
Full slow axonal transport model (kinetic diagram for this model is shown in Fig. 2b). (a) Concentration of cargos in the anterograde pausing state. (b) Concentration of cargos in the retrograde pausing state.

**Fig. S3.**
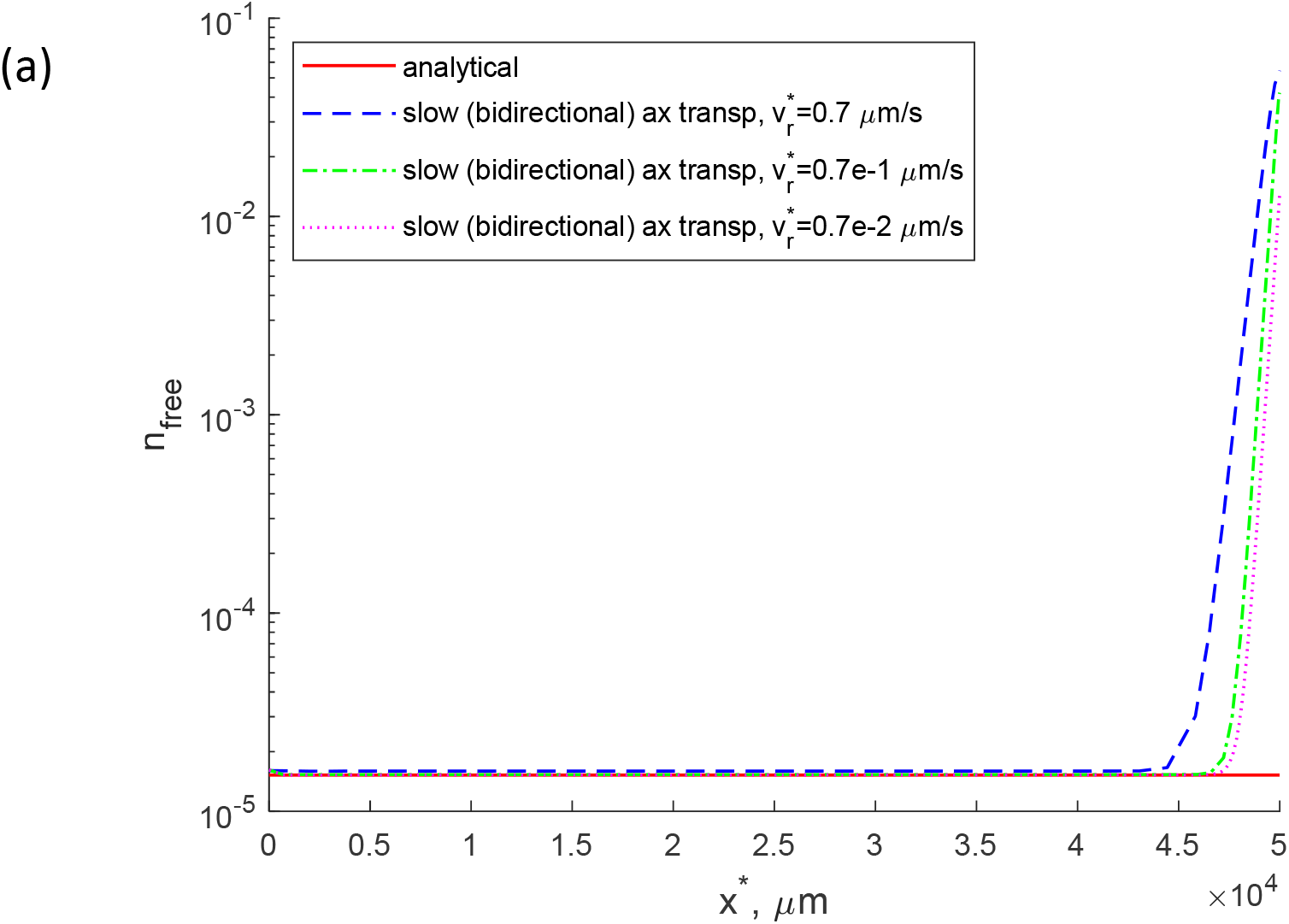
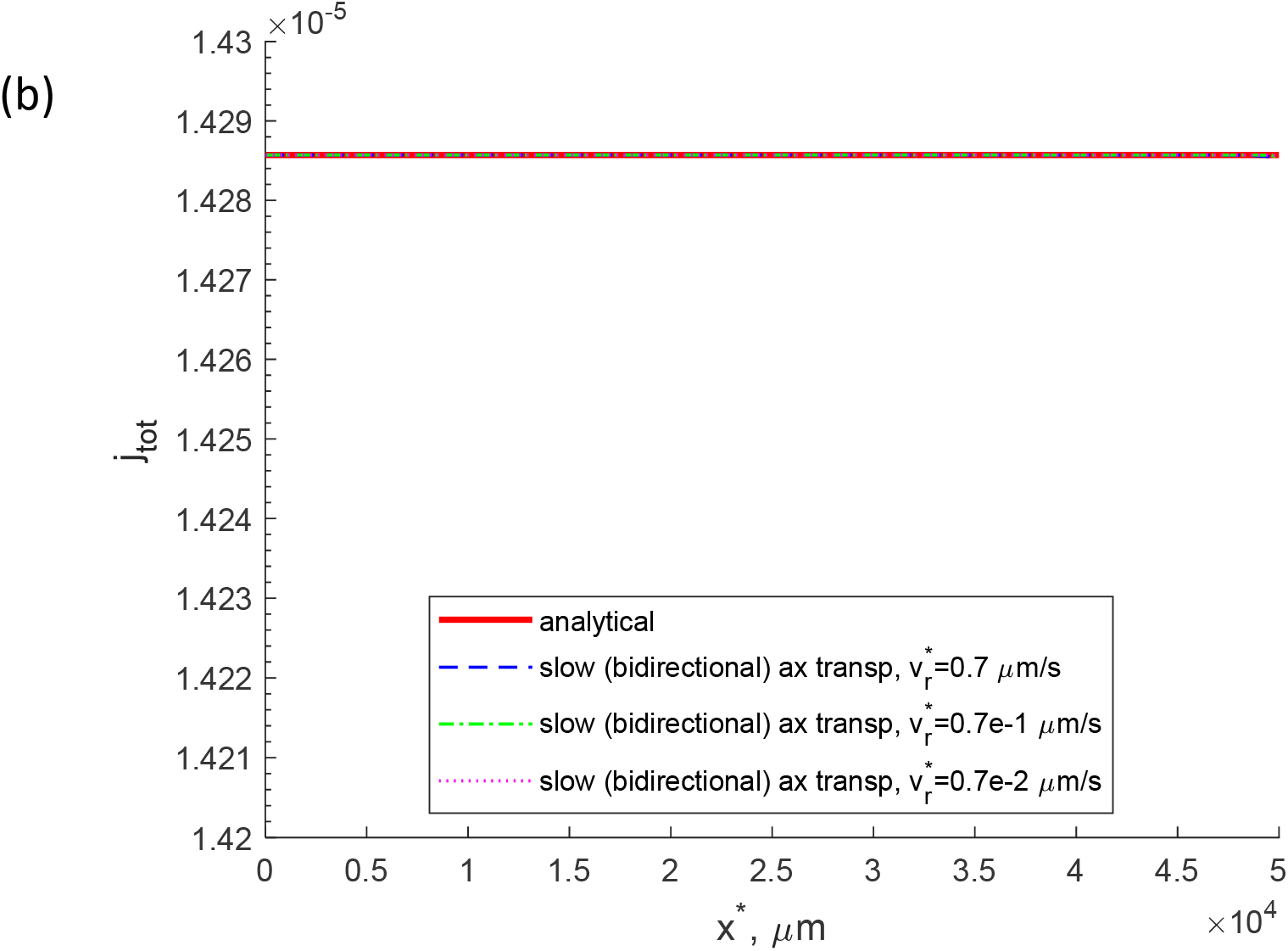
Full slow axonal transport model (kinetic diagram for this model is shown in Fig. 2b). (a) Concentration of cargos in the free (cytosolic) state. (b) Total flux of cargos due to the action of molecular motors and diffusion. 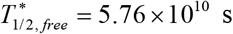.

**Fig. S4.**
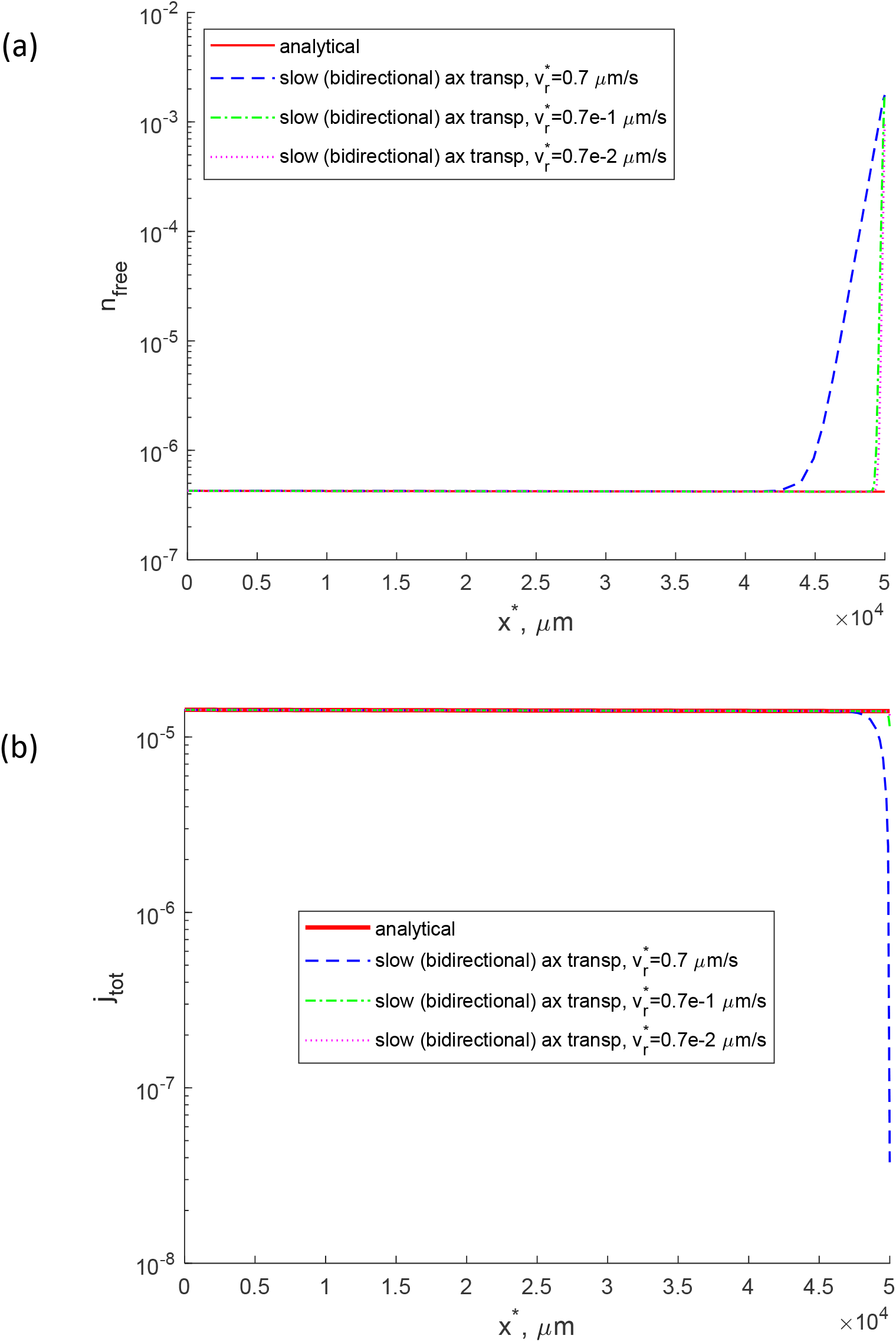
Full slow axonal transport model (kinetic diagram for this model is shown in Fig. 2b). (a) Concentration of cargos in the free (cytosolic) state. (b) Total flux of cargos due to the action of molecular motors and diffusion. 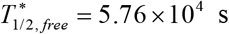.

